# Rampant prophage movement among transient competitors drives rapid adaptation during infection

**DOI:** 10.1101/2021.02.01.429245

**Authors:** Christopher W. Marshall, Erin S. Gloag, Christina Lim, Daniel J. Wozniak, Vaughn S. Cooper

## Abstract

Interactions between bacteria, their close competitors, and viral parasites are common in infections but understanding of these eco-evolutionary dynamics is limited. Most examples of adaptations caused by phage lysogeny are through the acquisition of new genes. However, integrated prophages can also insert into functional genes and impart a fitness benefit by disrupting their expression, a process called active lysogeny. Here, we show that active lysogeny can fuel rapid, parallel adaptations in establishing a chronic infection. These recombination events repeatedly disrupted genes encoding global regulators, leading to increased cyclic-di-GMP levels and elevated biofilm production. The implications of prophage-mediated adaptation are broad, as even transient members of microbial communities can alter the course of evolution and generate persistent phenotypes associated with poor clinical outcomes.

**One Sentence Summary:** Bacteriophage act as genetic regulators that are key to establishing chronic infections and are rapidly shared among co-infecting strains.

## Introduction

“A non-specific parasite, to which partial immunity has been acquired, is a powerful natural weapon.” (Haldane 1949)(*1*)

“But this weapon, if captured by the victim and re-deployed, can engender victory.” (this study)

We often begin study and treatment of chronic bacterial infections much as we might start watching movies already an hour underway – we do our best to understand the current scene despite missing much of the character and plot development. By the time an infection is recognized, hundreds of pathogen generations may have passed, during which the population may have evolved along cryptic paths that cannot be retraced. Consequently, the earliest factors governing the fate of a new bacterial infection remain unclear (*2*).

Because many infections arise from a single clonal population, longitudinal isolates from patients have yielded invaluable insight into how individual bacteria initially adapt and then persist in infections (*3–9*). By studying a time series of isolates or populations, the selective pressures experienced by pathogens can often be inferred retrospectively by determining what genetic and phenotypic traits were selected in the infecting population (*10–13*). What many of these studies missed, however, were the initial selective pressures and eco-evolutionary interactions between the pathogen and close competitors as the microbial community is established. If we were instead able to watch the movie from the beginning, we can understand why certain pathogens outcompete others and become successful colonizers in the new host.

One potential axis of competition between bacterial lineages is lysogenic bacteriophage that reactivate to become lytic (*14–19*). It has long been appreciated that interactions among competing species can be mediated by their shared predators or parasites (*20*), a process known as apparent competition. In the case of integrated prophage, reactivation during the stress of competition is often lethal to the host, but sibling clones with the dormant prophage remain resistant and benefit from the high mortality of susceptible competing populations lacking the phage. Thus, the fitness cost of bearing prophage can be overcome at a population level by attacks of the active phage against competitors.

The possibility that lysogenic phage may mediate fitness when initiating infection became evident in our recent study of *Pseudomonas aeruginosa* experimental chronic wound infections (*21*). Six different strains of *P. aeruginosa* were inoculated into a porcine full-thickness thermal injury wound model to determine which strain(s) were superior competitors and how they genetically adapted to the infection. The PA14 and PAO1 strains repeatedly prevailed in independent infections, and PA14 hyperbiofilm forming variants became detectable within days and persisted for weeks. These variants were caused by single mutations in the *wsp* pathway that induced hyperbiofilm formation by increasing levels of the key signal cyclic-di-GMP (*22*). Remarkably, some variants also gained CRISPR spacer insertions that conferred immunity to a temperate phage present in a co-inoculated strain (*21*). Therefore, we hypothesized that prophages mediate competition by killing sensitive competitors, selecting for resistant lineages, and generating new mutant genotypes that may be adaptive in the host environment. Here, we find that the genomes of winning clones contain multiple mutations caused by these lysogenic phages and other mobile genetic elements. In many cases, prophages were the sole means of adaptation over the course of the infection, and their speed and multiplicity of actions revise our understanding of the genetics of rapid evolution during infection.

## Results

### *P. aeruginosa* RSCVs are selected in porcine chronic wound infections

Given a mixture of bacterial strains added in equal fractions to a defined environment, picking the winner remains a daunting challenge. To determine which *P. aeruginosa* strain was competitively superior in a chronic infection, we used a porcine full-thickness thermal injury wound model that was co-inoculated with six different strains of *P. aeruginosa* and followed the infection for 28 days (Figure 1A). We previously reported the general outcomes of the infection (*21*), but describe some of the relevant data here for clarity. The starting inoculum was 10^8^ CFU/wound, with an equal starting distribution of *P. aeruginosa* strains PA14-1 and PAO1-B11 (burn wound isolates and model organisms), B23-2 (wound isolate), CF18-1 (non-mucoid cystic fibrosis isolate), MSH10-2 (water isolate), and S54485-1 (urinary tract infection isolate). By day 3 and continuing throughout the duration of the experiment, strains PA14 and PAO1 became predominant, while the other strains remained at or below the detection limit (<0.1%), based on neutral genetic markers. We used heritable differences in colony morphologies to screen for evolved variants over the course of the infection. The primary novel phenotype was the rugose small colony variant (RSCV), which accounted for up to 2% of the total *P. aeruginosa* population screened. The RSCV phenotype is well known to arise in chronic infections (*23–25*) and have a hyperbiofilm phenotype due to the overproduction of exopolysaccharides Pel and Psl (*22, 25, 26*) and the protein adhesin CdrA (*27*). Importantly, RSCVs are associated with persistence and worse clinical outcomes (*26, 28*). Following evolution in the wound environment, we sequenced both RSCV and non-RSCV isolates to identify the causative mutations of the RSCV phenotype. We used the standard approach of variant calling by mapping short, accurate reads (Illumina) to the ancestral genome. All of the RSCVs derived from PA14 had one of two mutations in the *wsp* operon (*21*), an important genetic locus for cyclic-di-GMP production and biofilm regulation in *P. aeruginosa* (*22, 29*). Surprisingly, however, no single nucleotide polymorphisms (SNPs) or short insertion/deletion (indel) mutations were detected among the majority of PAO1-derived RSCV isolates (12/17 sequenced RSCVs had no mutations; Table S1) or in most (16/28) PAO1 non-RSCV isolates sequenced (Table S2).

**Figure 1:**
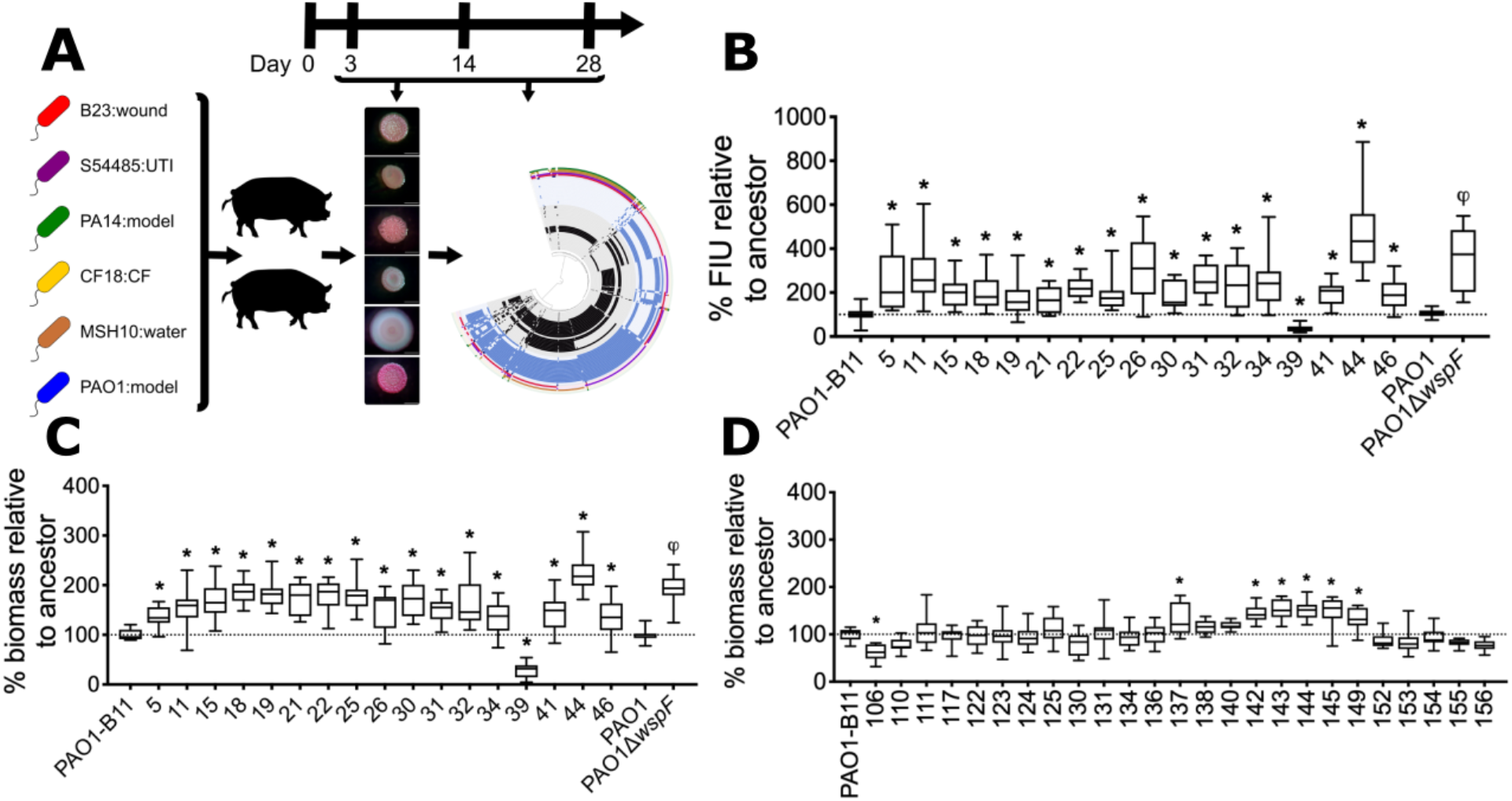
PAO1 RSCVs isolated from porcine burn wounds have a hyperbiofilm phenotype. **(A)** Summary of the experimental design. Six *P. aeruginosa* strains were inoculated into twelve wounds on two pigs and biopsies taken on days 3, 14, and 28 post infection. RSCVs were isolated and subjected to sequencing and analysis. **(B)** A cyclic-di-GMP reporter was electroporated into PAO1 RSCVs that were selected for sequencing. Cyclic-di-GMP levels were measured as green fluorescence and reported as GFP fluorescence intensity units (FIU) normalized to the ancestor strain, which was set at 100%. FIU was normalized to optical density. N=3, each with four technical replicates. Biofilms of PAO1 **(C)** RSCVs and **(D)** non-RSCVs that were selected for sequencing were grown in a 96 well plate for 4 h. Biofilm biomass was quantified by crystal violet. Biofilm biomass was expressed as a percentage, relative to the ancestor strain, which was set to 100 N=4, each with four technical replicates.%. * represents a p-value < 0.05 compared to the PAO1-B11 ancestor (one-way ANOVA with Tukey’s post-hoc test), and phi (φ) represents a p-value < 0.05 compared to PAO1 (Student’s t-test). Numbers on x-axis denote RSCV isolates.

### PAO1 RSCVs acquire adaptive mutations that increase biofilm production

Of the five PAO1 RSCV isolates with detectable mutations, only two (RSCV-41 and RSCV-44) had mutations in genes previously implicated in the RSCV phenotype. RSCV-44 had a single nucleotide polymorphism (SNP) in *wspF*(TàG, I68S) and RSCV-41 had a SNP in *retS* (AàC, T443P) (Table S1). RetS, the <u>r</u>egulator of <u>e</u>xopolysaccharide and <u>t</u>ype III <u>s</u>ecretion, is a sensor kinase that regulates the switch between acute and chronic infection, and when inactivated stimulates biofilm production through elevated cyclic-di-GMP levels (*30–32*). Introduction of *wspF*and *retS* wild type alleles *in trans* complemented the RSCV colony phenotype of these isolates, respectively (Figure S1A). This confirms that the SNPs in *wspF* in RSCV-44 and *retS* in RSCV-41 were responsible for the RSCV phenotype of these two isolates. The remaining three isolates with identified mutations (RSCV-11, RSCV-25 and RSCV-32; Table S1), had SNPs in genes hypothesized to be unrelated to the RSCV phenotype, as introduction of the wild type allele did not complement the variant colony phenotype (Figure S1B). That left 15 of 17 sequenced RSCVs with no obvious genetic cause for the RSCV phenotype.

The absence of SNPs or short indels in the PAO1 RSCV isolates was perplexing, given their heritable RSCV phenotypes (Figure 1A). Using the same approach, we previously identified mutational parallelism in the *wsp* operon in RSCVs derived from strain PA14 (*21*). Further, we confirmed that the heritable variant colony morphologies of PAO1 derived RSCVs was accompanied by elevated levels of cyclic-di-GMP and significant hyperbiofilm formation compared to the ancestor (Figure 1B and C). In contrast, the majority (22 of 28) of the PAO1 derived non-RSCVs recovered over the course of the infection did not have significantly elevated levels of biofilm production (Figure 1D) and no identifiable non-synonymous mutations were found in 21 of 28 non-RSCV isolates (Table S2).

### Newly acquired prophages are responsible for the RSCV phenotype

Reference-based mutation calling is largely blind to horizontal gene transfer (HGT) because any sequences not in the reference genome are unmapped and are not reported. We therefore hypothesized that the evolved PAO1 RSCV phenotypes resulted from HGT events between coinoculated strains. To discover possible HGT events we aligned each reference and isolate genome to one another and catalogued the presence or absence of homologous genomic regions. We discovered fragments of exogenous DNA, ranging from 7-70 kilobases, incorporated into nearly all of the PAO1 genomes isolated from the porcine wounds, both RSCV and non-RSCV (Figure 2). These foreign DNA sequences all mapped to mobile genetic elements, primarily prophages, found in one of the four other co-inoculated *P. aeruginosa* strains that eventually became undetectable in the wounds (Table S3). The amplification and mobilization of prophages, along with the demise of their donor genotypes, shows that transiently colonizing strains can substantially alter the genomes and frequencies of numerically dominant strains.

**Figure 2.**
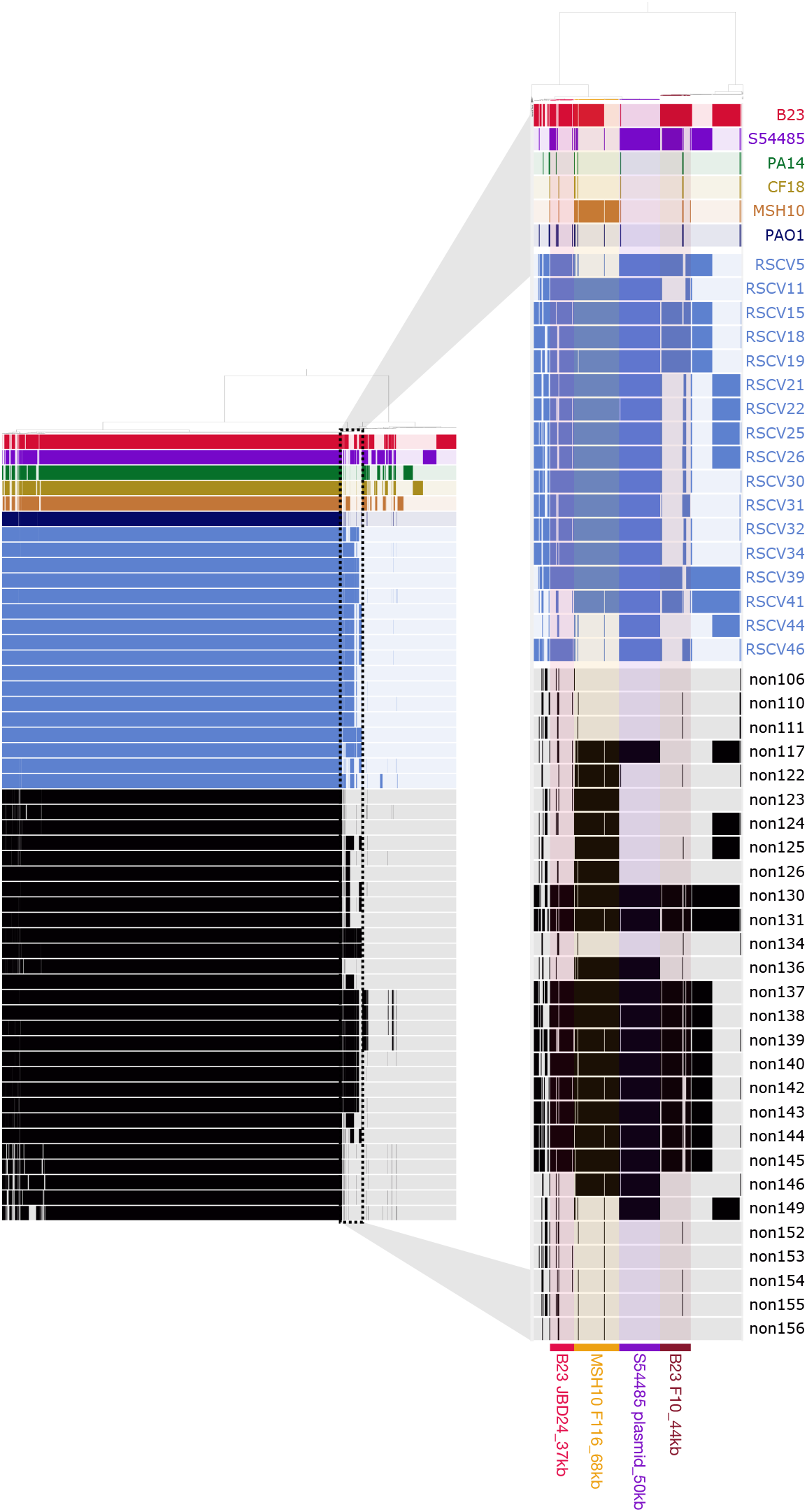
Genome alignments highlight the acquisition of mobile genetic elements by PAO1 isolates. Map of the pangenome containing the ancestral co-inoculated *P. aeruginosa* strains and the derived PAO1 isolates sequenced. Black bars indicate genome sequences belonging to the non-RSCV isolates, and blue bars indicate sequences belonging to the RSCV isolates. The six co-inoculated strains are in red (B23), purple (S54485), green (PA14), yellow (CF18), orange (MSH10) and dark blue (PAO1). The solid block of bars to the left represents the core *P. aeruginosa* genome shared by all strains. The zoomed-in area to the right represents part of the accessory genome that was largely absent in the PAO1 ancestor and acquired during evolution in the porcine wound. Presence of these bars in the accessory genome of the recovered PAO1 isolates indicate recently acquired mobile genetic elements.

The newly integrated prophages in PAO1 isolates originated from either the co-inoculated B23 or MSH10 strains. By day 3, these donor strains fell to at or below the detection limit from amplicon sequencing and were never detected by culturing methods (*21*), suggesting that prophage were activated and lysogenized the PAO1 population early in the infection. The prophage integrated in MSH10 is closely related to the F116 (*Podoviridae*) phage (*33*), is one of two predicted prophages in its genome (Table S4), and was found in 82% of sequenced RSCVs and 64% of non-RSCVs (Table S3). In addition, a prophage closely related to the Mu-like JBD24 (*Siphoviridae*) phage (*34, 35*) was identified in the B23 genome (Table S4) and was detected in 100% of the RSCVs and 50% of non-RSCVs (Table S3).

We predicted that these newly acquired prophage contributed to the RSCV phenotype. Through long read sequencing, we determined that many prophage genes, particularly from phage JBD24, repeatedly inserted into loci that could explain the RSCV phenotype (Table 1). Mutations (phage insertions and SNPs) in *retS* (PA4856) were found in the majority of RSCVs sequenced (10 of 17) and consisted of at least four separate mutational events across two animals and three different wounds (Table S5). All phage insertions into *retS* appeared to originate from the co-inoculated *P. aeruginosa* strain B23 (Table 1). This parallelism of HGT events at the gene level indicates positive selection on the resulting phenotype.

**Table 1.**
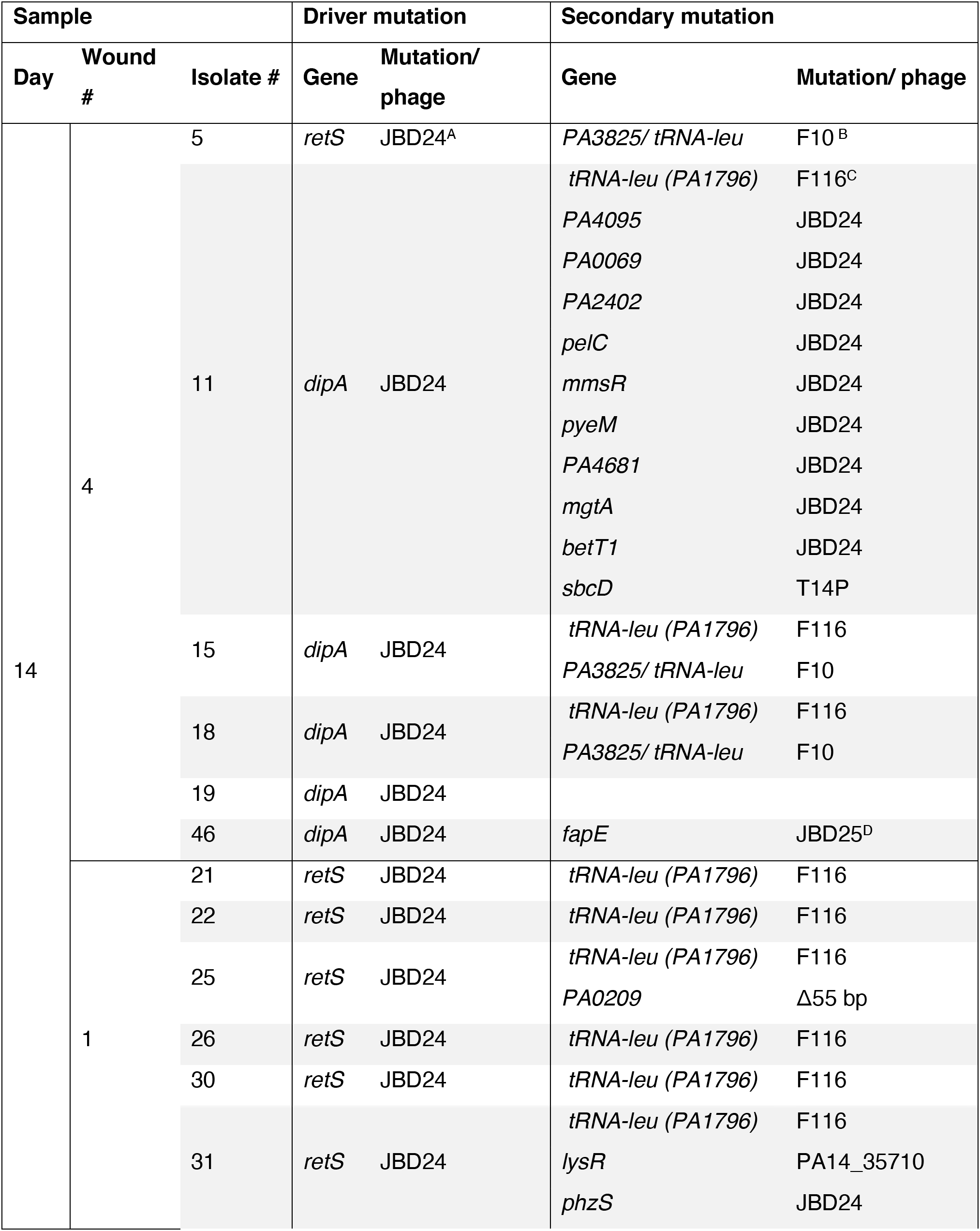
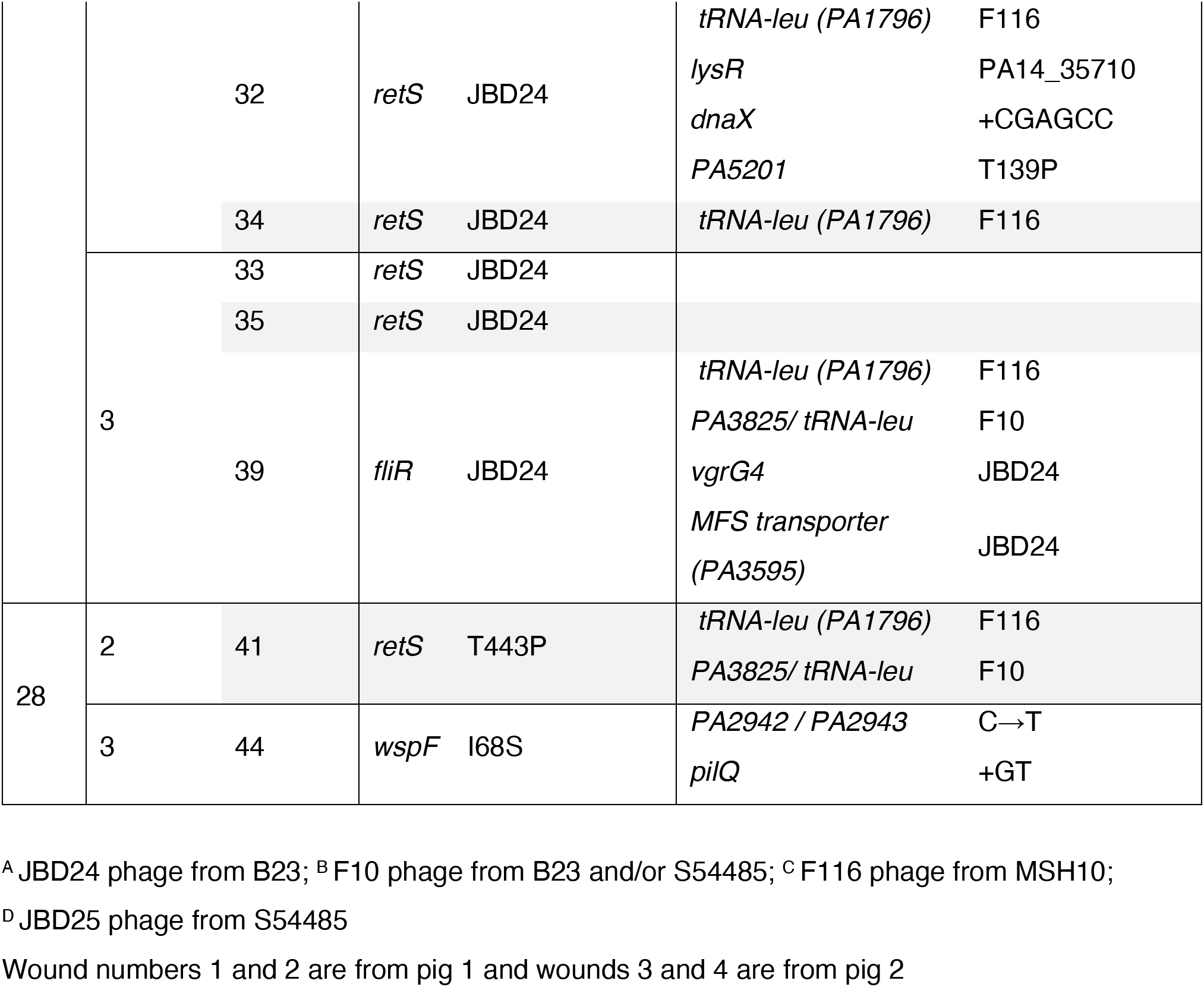
Mutations and mobile genetic elements in PAO1 RSCV isolates.

To test if phage-mediated inactivation of *retS* caused the RSCV phenotype, rather than by producing a novel phenotype from a phage-encoded gene, we introduced the wild type *retS* allele *in trans*. This restored the RSCV colony phenotype to the ancestral morphology (Figure 3). However, introduction of wild type *retS in trans* had no effect on the ancestral PAO1 strain or on an engineered *wspF* deletion mutant (PAO1Δ*wspF*) (Figure S2), demonstrating allele specificity. Therefore, loss-of-function *retS* mutations that derepress a signalling cascade and elevate cyclic-di-GMP production are hypothesized to be an important early adaptation in wound infections.

**Figure 3.**
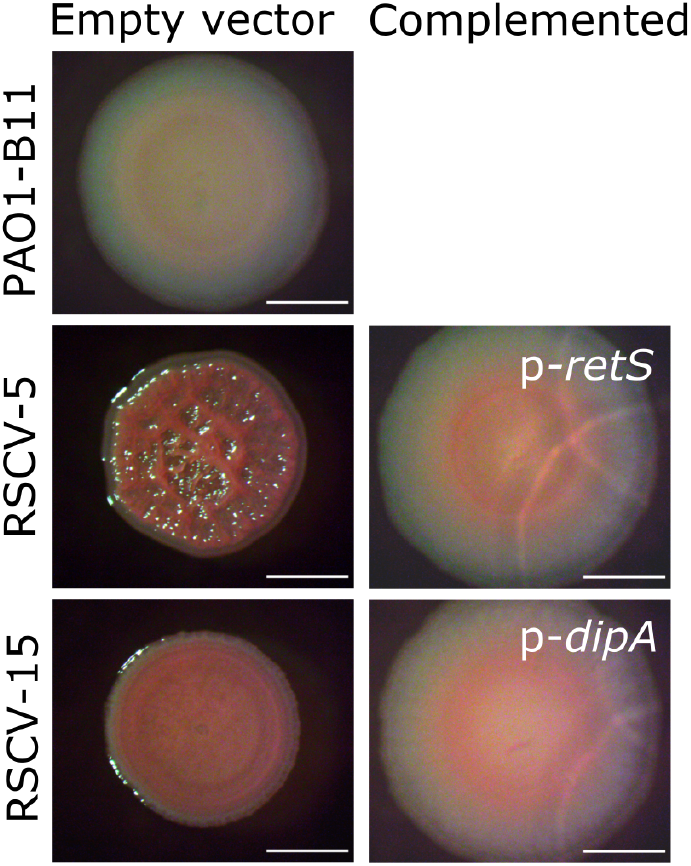
Disruption of genes involved in cyclic-di-GMP regulation by prophage insertion is responsible for the PAO1 RSCV phenotype. Representative PAO1 RSCVs were complemented by introducing the wild type copy of the gene disrupted by prophage insertion. Parent strains (empty vector; pUCP18) and complemented strains were grown on VBMM and colony morphology assessed. RSCV-5 was selected as a representative *retS*-disrupted RSCV, and RSCV-15 was selected as a representative *dipA-*disrupted RSCV. PAO1-B11 is the ancestor strain that the RSCVs evolved from. Scale bar indicates 2mm.

In addition to the recurring *retS* mutations, phage insertions into other genes responsible for the RSCV phenotype were observed. Five isolates (RSCV-11, −15, −18, −19 and −46; all from the same wound) gained a prophage insertion in *dipA* (PA5017) (Table 1). DipA, dispersion-induced phosphodiesterase <u>A</u>, is responsible for initiating biofilm dispersion. DipA deletion mutants have increased exopolysaccharide production, dampened swimming and swarming motility, and enhanced initial attachment (*36, 37*). Complementation of RSCV-15 and −46 with the wild type *dipA* allele *in trans* restored the colony morphology to the ancestral type, demonstrating that prophage insertions in this gene caused the RSCV phenotype (Figure 3 and S1A). Here again, the prophage-mediated adaptation was not caused by the addition of new genes, but rather by disruption of an existing regulatory gene, which is a common initial route to increased fitness in new environments (*38–40*). Further, competition between strains in chronic wounds can generate eco-evolutionary dynamics, where induced prophage can infect and kill susceptible competitors and favor resistant types, but they can also produce new genotypes that are adapted to the phage-laden environment.

Mutations in *dipA, retS*, and *wspF* explained the phenotype of 16 of 17 PAO1 RSCVs, leaving only one unexplained genotype-phenotype link. This remaining isolate, RSCV-39, had the variant colony phenotype but was significantly impaired in biofilm production (Figure 1B) and had reduced levels of cyclic-di-GMP (Figure 1C). This isolate had a prophage insertion in *fliR* (Table 1), which is involved in flagella biosynthesis. A recent study linked mutations in flagellar biosynthesis pathways to the RSCV phenotype, but only when the mutants were grown on a solid surface (*41*). We therefore repeated the cyclic-di-GMP quantification of RSCV-39 by comparing levels in planktonic and surface grown cells. Consistent with the previous study (*41*), RSCV-39 only displayed elevated levels of cyclic-di-GMP when grown on a surface (Figure S3). These results, together with previous observations of flagella-mediated RSCVs, suggests that the disruption of *fliR* by prophage insertion is responsible for the RSCV phenotype in RSCV-39.

Interestingly, all of the sequenced PAO1 RSCV isolates, and 50% of the PAO1 non-RSCV isolates, also acquired a putative conjugative plasmid from the strain S54485. This plasmid had not been annotated in the S54485 genome, but circularized in our assemblies and shares homology with plasmid sequences from *Burkholderia pseudomallei* (CP009154) (*42*) and an *Acinetobacter baumannii* clinical isolate from Thailand (ERS1930304). The vast majority of the predicted proteins encoded on this plasmid were annotated as hypothetical and the role that this plasmid may play in host adaptation is under further investigation.

### Acquisition of new prophages provides immunity to phage re-infection

Given that prophages can provide resistance to phage superinfection (*43*), we hypothesized that the newly acquired prophages in the PAO1 wound isolates could provide immunity to phages isolated from the co-inoculated *P. aeruginosa* strains. To test this hypothesis, we isolated the JBD24-like and F116-like phages from strains B23 and MSH10, respectively, since these were most commonly integrated into the PAO1 isolates. Phages isolated from both strains were able to inhibit the growth of the ancestral PAO1 strain (Figure 4, Figure S4A) and the engineered PAO1Δ*wspF* mutant (Figure S5). However, the growth of RSCV-5 and −44, containing mobile elements from B23 and S54485, and RSCV-15 and −41, containing mobile elements from B23, MSH10, and S54485 (Figure 2), were unaffected by the isolated phages, demonstrating acquired immunity (Figure 5A and S6B-E). More noteworthy, RSCV-5 and −44 did not acquire genetic elements from MSH10 (Figure 2) but nonetheless became resistant to MSH10-derived phages (Figure 4A), suggesting cross-resistance. However, unlike RSCVs, non-RSCVs display immunity to the isolated temperate phages in a lysogen-dependent manner (Figure 4B and S4F-H).

**Figure 4.**
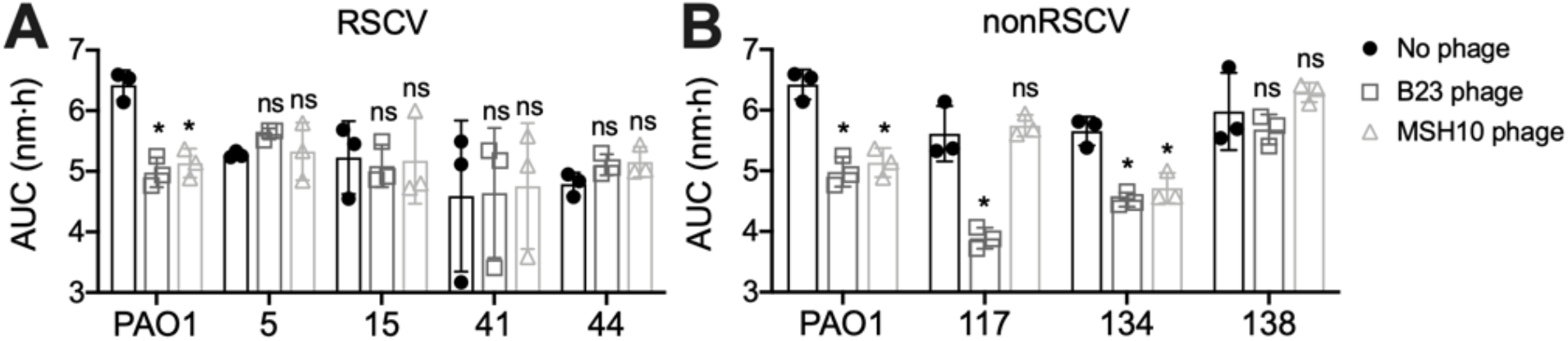
PAO1 wound isolates are immune to phage infection. Representative PAO1 **(A)** RSCV and **(B)** nonRSCV wound isolates were grown in planktonic culture with phage isolated from B23 (open squares) or MSH10 (open triangles) for 16 h and the OD measured every 30 min. Data is presented as area under the curve (AUC) of the growth curves depicted in Figure S7. * indicates p<0.05 compared to no phage control (solid circle) (one-way ANOVA with a Tukey’s post-hoc test); ns indicates no significant difference. N=3, each with three technical replicates. Data presented as mean ± SD, with the individual data points reflecting the mean of each biological replicate.

**Figure 5.**
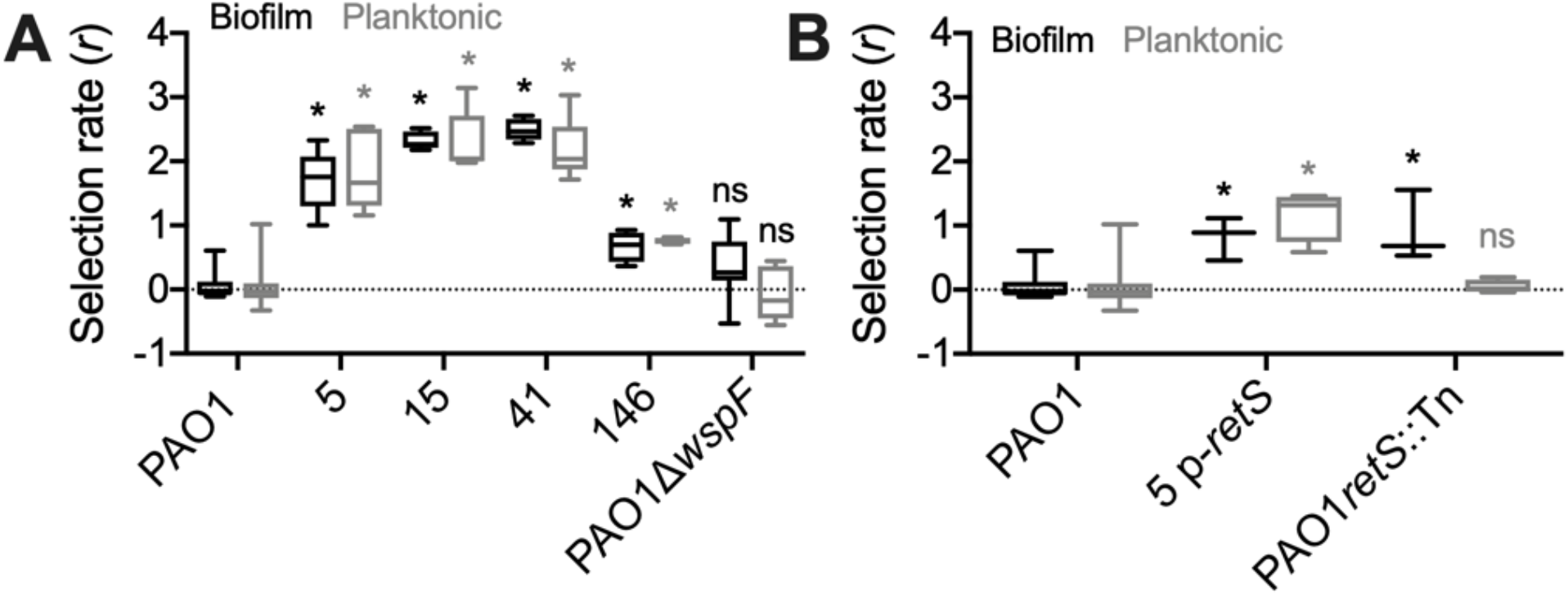
PAO1 RSCVs have increased fitness relative to the ancestral PAO1. PAO1 strains were complemented against the ancestral parent tagged with *lacZ* for 48 h in both biofilm (black) and planktonic (grey) conditions for 48 h. Fitness of the competing strain was determined by calculating the selection rate (r). **(A)** Fitness of representative PAO1 wound isolates relative to the ancestral PAO1 strain. PAO1Δ*wspF* competed against the isogenic PAO1 parent was used as a representative *wsp* mutant, instead of RSCV-44 due to the secondary SNP in *pilQ* in this isolate (Table S1). N=5. **(B)** To separate the contributions of the *retS* mutation and the presence of newly acquired prophage to the increased fitness phenotype, RSCV-5 containing the *retS* complementing plasmid (p-*retS*), and a PAO1 *retS* transposon mutant were completed against their parent PAO1 strains. N=3. * indicates p<0.05 compared to the PAO1 pairwise competition (one-way ANOVA with Tukey’s post-hoc test); ns indicates no significant difference.

### Extreme fitness advantages of evolved RSCV-causing mutations

Having established that activated prophage both selected and produced the isolated variants, we asked if fitness effects of lysogeny extended beyond resistance to superinfection. The regulatory genes disrupted by prophage alter phenotypes like RSCV, but new genes encoded by the prophage could also influence fitness. Measuring fitness, even *in vitro*, can be predictive of pathogen success in the host (*44*). Isolates were competed against the *lacZ*-marked ancestral strain over 48 hours in biofilm and planktonic conditions. Both types of *retS* mutants, whether the T443P substitution (RSCV-41) or a prophage insertion mutant (RSCV-5), were significantly more fit, outcompeting the PAO1 ancestor in both planktonic and biofilm conditions (Figure 5A). This was also the case for the *dipA* prophage insertion mutant (RSCV-15) (Figure 5A).

To distinguish effects of *retS* mutations from effects of prophage acquisition, we competed PAO1 against a transposon mutant of *retS* (PAO1*retS*::Tn; (*45*)) and against RSCV-5 complemented with wild type *retS* allele *in trans* (RSCV-5 p-*retS*). PAO1*retS*::Tn was more fit than the ancestor only in biofilm conditions; however, RSCV-5 p-*retS* was significantly more fit in both planktonic and biofilm conditions, which demonstrates benefits of lysogeny beyond disrupting *retS* (Figure 5B). Furthermore, a mutant that acquired several mobile genetic elements, but no observable gene disruptions or variant phenotypes (non-RSCV-146, Table S6), outcompeted PAO1 (Figure 5A), although to a lesser extent than RSCV-5 (Figure S7). Therefore, the enhanced competitive fitness of PAO1 RSCVs compared to the ancestor appear to result from combined effects of the RSCV hyperbiofilm phenotype and the horizontally acquired mobile genetic elements themselves (Figure 5B).

Given the focused selection on *wsp* mutants of PA14 (*21*), we were surprised that homologous mutants were not found in PAO1. We therefore competed PAO1Δ*wspF* against the isogenic PAO1 parent, in both planktonic and biofilm conditions. Contrary to the high fitness advantage of Δ*wspF* observed in PA14 (*21*), PAO1Δ*wspF* exhibited no fitness advantage, even in biofilm growth conditions (Figure 5A). These results contrast with the high selective advantage of *retS* and *dipA* mutants had over the PAO1 ancestor (Figure 5). Further, in direct pairwise competition, RSCV-5 (the *retS* prophage insertion mutant) was significantly more fit than PAO1Δ*wspF* (Figure S7B), again suggesting why PAO1 *retS* mutants may be selected *in vivo* over PAO1 *wsp* mutants.

The relative fitness values of the evolved RSCV isolates indicate extreme competitive advantages that could indicate active exclusion or killing of the competitor. Specifically, the ancestral PAO1 strain was rapidly outcompeted and became undetectable after 24-48 hours in many of these fitness assays. We also observed zones of inhibitions around the RSCV isolates when plated adjacent to the ancestor (Figure S8), suggesting active killing of the ancestral strain by the RSCV isolates. To test this hypothesis, we plated colonies of evolved RSCV isolates next to the ancestor and filmed their interactions while growing over time (Supplemental Movies 1-4). These movies indicated that both the *retS* T443P SNP (RSCV-41) and the *retS* prophage insertion mutant (RSCV-5) outcompeted the ancestral strain by direct inhibition. The inhibition by the insertion mutant, RSCV-5, appears to be through phage lysis. In support of this, we found that this newly acquired prophage in RSCV-5 can be induced to kill the ancestral strain (Figure S5). However, the mechanisms of the observed direct inhibition are still under investigation.

## Discussion

For any given set of bacterial strains, the fittest in a given environment is usually unpredictable, regardless of available data. But a growing literature shows that bacterial fitness is often both at the mercy of, and empowered by, prophage (*46, 47*). These enemy cargo harm the carrier when activated but can kill non-lysogenized competitors, providing significant competitive advantages to resistant genotypes. In this study, when mixed populations of *P. aeruginosa* strains were added to model chronic wounds, the earliest dynamics and adaptations occurred in response to prophage, but not in the typical manner where lysogens prevail because they resist superinfection. Rather, the four strains that contributed at least four mobile elements to this strain mixture (Table S4, Figure 2) were swiftly outcompeted, as these elements selected for CRISPR-mediated resistance in some clones of PA14 (*21*), and produced an assortment of new lysogens in the PAO1 background. Newly lysogenized PAO1 variants also acquired novel adaptations to the wound environment beyond phage resistance, including hyperbiofilm traits that may enhance resistance to host defenses (*28*). The ecological theory of apparent competition can be summarized as “the enemy of your enemy is your friend”; for example, when one host species carrying a pathogen that it evolved to tolerate outcompetes another species that is more susceptible. The PAO1 variants characterized here upend this dynamic by capturing prophages that generate adaptations via the genes they disrupt and the defenses to superinfection that they encode.

Here we demonstrate that transient ecological interactions alter the course of an infection through prophage movement and subsequent selection on prophage-disrupted genetic targets. Prophages that emerged from strains that were rapidly outcompeted provided substantial fitness benefits to the winning strains and increased the mutation supply upon which selection could act. We identify the causes of at least part of that enhanced competitiveness as effects of the genetic loci disrupted by inserted prophage. PAO1-derived RSCVs with prophage insertions in *retS* were routinely isolated from the porcine chronic wounds (Table 1). By disrupting the RetS signaling cascade that regulates biofilm formation and virulence (*30, 31*), the RSCVs could outcompete other ancestral strains in the wound environment. This discovery has two major consequences: (1) short term exposure to closely related strains can alter the evolutionary trajectory of bacterial populations, and (2) selection for hyperbiofilm formation is almost certainly adaptive in an infection, as has been previously hypothesized (*21, 25, 48, 49*).

Many chronic infections begin clonally and temporally diversify as the infecting strain adapts to new niches in the host (*3, 4, 50–52*). This has been elegantly demonstrated for cystic fibrosis lung isolates, where the phylogenies indicate a single originating colonizing strain giving rise to multiple coexisting lineages (*6, 10, 50*). However, our findings here suggest an alternative outcome: several competing strains may infect a host and dictate the evolutionary fate of the predominant strain. In this model, the infection would still appear clonal because an invading or co-infecting strain would be outcompeted by the more fit, abundant strain (a process that only takes a few days at most according to our results (*21*)). However, these transient co-infecting strains may significantly and cryptically alter the targets of selection. Given that patients are exposed to many opportunistic pathogens in the environment and in clinical settings during treatment (*53–58*), this scenario seems possible and still consistent with the ultimately clonal appearance of chronic infections. An additional piece of evidence supporting our hypothesis is that most of the sequenced isolates from chronic infections have many genomic regions that appear to be recently acquired mobile genetic elements, including phage (*19, 59*). This is usually thought to be from competition in the environmental reservoir. This still may be the case, but it could also be due to previous competition within the host that drove early adaptations in the predominant strain.

Prophages are often maintained in the bacterial chromosome because they provide resistance to phage superinfection (*43*), but they can also enhance competitiveness in many environments including *in vivo* (*59*). In contrast to the better understood path of lysogenic conversion where the bacterium gains a fitness advantage from a horizontally acquired gene, active lysogeny involves prophage insertion into a gene or regulatory region that inactivates that gene, creating a sort of genetic switch (*60*). In this study, active lysogeny disrupted different regulators of biofilm formation and improved fitness *in vitro* and *in vivo*. Mutations in *retS* that increase biofilm formation and type VI secretion are thought to be adaptive in chronic infections (*9, 13*), as this signaling cascade is a central switch between acute and chronic infection (*30, 31*). Other examples of active lysogeny include prophages causing mutator phenotypes in *Streptococcus pyogenes* by disrupting *mutL* (*61, 62*), prophages disrupting *com* genes required for *Listeria* virulence and survival in phagosomes (*47, 63*), and the classic example of prophage regulating sporulation cycle components (*64–68*). Given the potentially reversible nature of active lysogeny and technical difficulty in observing these events, it may be that active lysogeny is more common than previously appreciated, especially in environments where competition among related strains is prevalent. Because active lysogeny contributes to a growing list of pathogenic traits, from virulence to sporulation and now, hyperbiofilm formation, the mechanisms and scope of active lysogeny demand further inquiry.

There is a growing body of work demonstrating that mobile genetic elements are a significant source of mutation driving increased biofilm formation in infections. In a patient with chronic *P. aeruginosa* pulmonary disease, insertion sequences of the ISL3-family disrupted genes involved in flagellar function, type IV pili, and other genetic loci that increased virulence of the mutants (*69*). Davies *et al*. observed *in vitro* a similar mechanism of phage-mediated adaptation to what we report here *in vivo* (*70*). In that study, an evolution experiment with PAO1 was conducted with added temperate phages, some of which inserted in targets relating to biofilm formation or motility, including *dipA* (*70*). Further, it has long been known that adding phages to biofilms can select for small-colony variants and that active phages have been observed in many chronic infection isolates (*71–73*). Our work synthesizes these findings by showing how competing strains can provide a source of the infecting phages, which in turn produce newly infected genotypes that are selected *in vivo* to enhance biofilm production and maintain the prophage in the population.

Two main takeaways emerge that advance our understanding of the evolutionary forces and selected traits within opportunistic infections. The first is that mutations that increase cyclic-di-GMP levels and therefore increase biofilm formation are adaptive in wounds. This result adds to a growing body of evidence from patients that biofilm formation is adaptive in chronic infections (*25, 49*). Our study shows that different strains of *P. aeruginosa* may take different routes to higher cyclic-di-GMP levels, where the convergent evolution of a hyperbiofilm phenotype may be caused by mutations in *wsp*, *retS*, *dipA*, or one of the other diguanylate cyclases or phosphodiesterases (*74–77*). The prevalence of these mutations that increase cyclic-di-GMP across many different hosts and studies strongly support that these are beneficial mutations, but rarely are they the most abundant mutation in the host infection (*21, 78*). An explanation for their prevalence but low abundance could be negative frequency dependent selection, where RSCVs are more fit when rare but disfavored at high frequency (*79, 80*). Nonetheless, the constellation of routes to RSCV’s and their relative rarity may make clinical targeting of this phenotype challenging. The second takeaway from this work is that inter-strain, and likely inter-species, interactions can significantly alter the evolutionary dynamics of an infection. Dormant or extinct strains that were initially present at equal ratios in the wounds provided prophage that infected and altered genes in the eventual winners, producing new genotypes and phenotypes for selection to act upon in addition to the hyperbiofilm phenotype.

Ultimately, the work presented here should inspire a broader view of how natural selection operates in an infection, where transient interactions in the microbial community and the contributions of mobile genetic elements can define the evolutionary-genetic course of an infection. These multi-strain or species interactions can happen early in the infection, in a window of time often missed by current surveillance methods. Therefore, catching the movie in the middle is not sufficient to understand how the characters behaved in the end. Instead, initial pathogen adaptations in chronic infections provide crucial plot development that needs to be examined more closely in order to better anticipate the rest of the story.

## Acknowledgments

Acknowledgments follow the references and notes but are not numbered. Acknowledgments should be gathered into a paragraph after the final numbered reference. This section should start by acknowledging non-author contributions and then should provide information under the following headings:

## Author contributions

CWM, ESG, DJW, and VSC conceptualized the study. CWM, ESG, and CL were responsible for data generation and data curation. CWM and ESG wrote the original drafts and all authors reviewed and edited subsequent writing;

## Data and materials availability

All data are included in results and supplementary materials except the following: all Illumina and Nanopore sequences have been deposited in the NCBI Sequence Read Archive (SRA) to the BioProject PRJNA633671 under accession numbers SAMN14968233 to SAMN14968292. Code for sequence processing can be found here: https://github.com/sirmicrobe/pig_wound_manuscripts/.

