## Supplementary Materials for "Rampant prophage movement among transient competitors drives rapid adaptation during infection"

Rampant horizontal gene transfer among transient competitors drives rapid adaptation during infection

**This PDF file includes:**

Materials and Methods
Figs. S1 to S8
Tables S1 to S7
Captions for Movies S1 to S6

**Other Supplementary Materials for this manuscript include the following:**

Movies S1 to S6

### Materials and Methods

#### *Porcine infection model*

As previously described (*1*), two pigs were each given six, two inch full-thickness third-degree thermal injury wounds that were covered with impermeable bandages. Three days after injury, each wound was inoculated with a total of $10^8$ CFU/mL of a *P. aeruginosa* consortium. The consortium consisted of six different strains of *P. aeruginosa*, including equal parts of PA14-1, PAO1-B11, B23-2 (wound isolate), CF18-1 (non-mucoid cystic fibrosis isolate GenBank ID; NZ_KI519281), MSH10-2 (water isolate GenBank ID; NZ_KE138672) and S54485-1 (urinary tract infection isolate GenBank ID; NZ_KI519256) that were tagged with a unique barcode at the *Tn7* site prior to infection (*1*).

#### *Isolating evolved variants from wounds*

On days 3, 14, and 28 post infection, replicate 8 mm punch biopsies were taken from two wounds on each pig. Biopsies were homogenized in PBS and plated on *Pseudomonas* isolation agar and Vogel-Bonner minimal media (0.2 g/L $MgSO_4 \cdot 7H_2O$, 3.5 g/L $NaNH_4HPO_4 \cdot 4H_2O$, 10 g/L $K_2HPO_4$, 0.1 g/L $CaCl_2$, 2g/L citric acid, 1 g/L casamino acid, 40 µg/mL Congo red, 15 µg/mL brilliant blue, solidified with 1% agar; VBMM), each supplemented with 100 µg/mL gentamicin. Heritable phenotypic variants were counted to determine the frequency of mutant phenotypes over the course of the infection. Isolates were maintained on either *Pseudomonas* isolation agar or lysogeny agar or broth (LA/B).

#### *Cyclic-di-GMP quantification*

The cyclic-di-GMP fluorescent reporter p*cdrA*::*gfp*, or empty vector pMH487 (*2*), was electroporated into PAO1 RSCVs and ancestral parent. Cultures were grown overnight in LB with 300µg/mL carbenicillin to maintain plasmid selection. Overnight cultures were diluted to an $OD_{600}$ 0.1. Cells were pelleted and resuspended in PBS. 100 µL was transferred to a black 96-well plate (Corning). $OD_{600}$ and green fluorescence were measured on a SpectraMax i3 plate reader (Molecular Devices). Green fluorescence values were normalized to OD and auto-fluorescence from the empty vector control was subtracted from each reading. Three biological replicates were performed, each with four technical replicates.

#### *Colony morphology complementation*

Desired genes were amplified by PCR using primers detailed in Table S8 and genomic DNA isolated from wild type PAO1. Primers were designed with restriction enzyme sites at the 5' end and are detailed in Table S7. PCR products were purified using the QIAquick PCR purification kit (Qiagen; 28106) according to manufactures protocol. Purified products and pUCP18 (*3*) were digested with appropriate restriction enzymes (New England Laboratories; NEB) according to manufactures protocol. Restriction enzymes were heat inactivated at 80°C for 15min before digested PCR products were ligated into linearized pUCP18 using T4 ligase (NEB) according to manufactures protocol. 5 µL of the ligation reaction was transformed into chemically competent *Escherichia coli* NEB5α cells and plated onto LA plates supplemented with 100 µg/mL ampicillin, and 100 μg/mL IPTG and 40 μg/mL X-gal for blue/white colony selection. Constructs were confirmed by sequencing. Confirmed constructs were purified using QIAprep Spin Miniprep kit (Qiagen; 27106) according to manufactures protocol. Constructs were electroporated into appropriate PAO1 RSCVs and the ancestor PAO1-B11 strain. To complement *wspF* mutations, pSP5 plasmid (*4*) was used.

To determine if introduction of the wild type gene *in trans* could complement the RSCV colony morphology, overnight cultures of appropriate strains were grown in LB supplemented with 300 μg/mL carbenicillin. 1 μL of the overnight culture was spotted onto VBMM plates supplemented with 300μg/mL carbenicillin and incubated overnight at 37°C. Colonies were imaged on a Stereo Microscope (AmScope) fitted with a Microscope Digital Color CMOS camera (AmScope). Images were processed in FIJI (*5*).

***Biofilm assay***

Overnight cultures were diluted to 0.5 $OD_{600nm}$ in VBMM broth and 100 μL transferred to a micro-titer 96-well plate. Biofilms were incubated in a humidified chamber at 37°C for 4 h. Biofilms were washed three times in PBS and remaining biomass stained with 120 μL 0.1% crystal violet for 30 min at room temperature. Biofilms were washed three times in PBS and crystal violet extracted in 150 μL ethanol for 30min at room temperature. Absorbance was then quantified on a plate reader at an $OD_{590nm}$. Four biological replicates were performed, each with four technical replicates.

***Fitness assays***

In order to determine fitness of the evolved variants, pairwise competitions were performed with at least three replicates each in biofilm and planktonic. We performed twelve competitions: 1. PAO1-B11 vs. $PAO1^+$ (*lac*-marked), 2. mPAO1 (control for transposon mutants) vs. $PAO1^+$, 3. PAO1pUCP18 (empty vector control for complemented strain) vs. $PAO1^+$, 4. Δ*wspF* vs. $PAO1^-$, 5. RSCV-41 (SNP in *retS*) vs. $PAO1^+$, 6. RSCV-5 (insertion in *retS*) vs. $PAO1^+$, 7. RSCV-5 vs. Δ*wspF*, and 8. RSCV-15 (insertion in *dipA*) vs. $PAO1^+$, 9. *retS*::Tn (transposon insertion in retS) vs. $PAO1^+$, 10. RSCV-5::*retS* (RSCV-5 complemented with ancestral *retS*) vs. $PAO1^+$, 11. non-RSCV-146 vs. $PAO1^+$, 12. RSCV-5 vs. non-RSCV-146. To start the competitions, each isolate was grown independently in 5 mL of tryptic soy broth overnight at 37°C and then mixed at 1:1 ratio in the competition tube containing 5 mL tryptic soy broth. For planktonic, the competitors grew for 24 h and then 50 μL were transferred to a new tube for another 24 h. For biofilm, cultures grew on a 7 mm polystyrene bead for 24 h and then the bead was transferred to a new tube with sterile beads and allowed to grow for another 24 h (*6*, *7*). At 24 h and 48 h timepoints, planktonic cultures were serially diluted and plated for colony forming units (CFUs) on 1.5% tryptic soy agar with 80 mg/mL X-Gal. At the same timepoints, biofilms were harvested by transferring a bead in 1 mL PBS and sonicated with a probe sonicator for 10s at 30% intensity (Fisher Scientific Model FB120). Dispersed biomass was serially diluted and plated for CFUs. From these counts the selection rate, r, was calculated as the ratio of the log-transformed growth of each competitor over 24 h (*8*).

#### *Phage susceptibility testing*

To induce and purify prophages from desired *P. aeruginosa* strains, an overnight culture was diluted 1:100 and supplemented with 0.5 µg/mL of mitomycin C. The culture was then incubated at 37ºC at 200 rpm for 4 h or until the culture reduced in $OD_{600}$ by half. The cells were then pelleted and the supernatant filter sterilized.

To determine the susceptibility of *P. aeruginosa* strains to isolated phages, both plaque and planktonic assays were used. For the plaque assay, 200 µL mid-log *P. aeruginosa* culture was incubated with 100 µL phage lysate for 15 min at 37ºC. This was then mixed with 5 mL molten soft agar (LB solidified with 0.7% agar) supplemented with 10 mM $CaCl_2$ and $MgSO_4$ and poured over solidified hard agar (LB solidified with 1.5% agar). Plates were incubated overnight at 37ºC. The number of plaques were counted and enumerated for plaque forming units (PFU)/mL. Plaque assays were performed in triplicate.

To obtain a higher phage titer required for the planktonic assay, plaques were cut out and incubated in 5 mL LB for 30 min. The tube was vortexed at 1min intervals. The phage suspension was filter-sterilized, and plaque assay repeated using PAO1 as the host strain to enumerate PFU/mL. For the planktonic assay, mid-log P. aeruginosa culture was diluted to $10^6$ CFU/mL and purified phage suspension was diluted to $10^{12}$ PFU/mL. To a 96-well plate, 50 µL of both bacteria and phage were added to individual wells, generating 100 µL infections, with a final multiplicity of infection (MOI) of $10^6$. This high MOI was required to more readily observe infection by phages isolated from B23. Bacterial culture with 50 µL LB media was used as a no phage control. The 96-well plate was covered with a breathable sterile adhesive membrane. $OD_{600}$ was measured on a SpectraMax i3 plate reader (Molecular Devices) every 30 min for 16 h at 37ºC. Prior to each reading the plate was shaken at medium speed for 10 s. Planktonic assays were performed in triplicate, each with triplicate technical replicates. Area under the curve (AUC) of the mean of each biological replicate was determined using the analysis function in GraphPad Prism v.8 (GraphPad Software).

#### *DNA extraction and sequencing analysis*

Genomic DNA was extracted from both phenotypic variants and colonies with the ancestral phenotype using the DNeasy Blood and Tissue Kit according to the manufacturer's protocol (Qiagen). DNA was prepared for sequencing using a modified Illumina Nextera protocol (*9*). DNA was sequenced with an Illumina NextSeq 500 at the Microbial Genomics Sequencing center (migscenter.com). Reads were quality filtered and trimmed with trimmomatic v0.36 (*10*) and then variants were called using *breseq* v0.30.0 or higher (*11*). Reads were mapped back to the PAO1 RefSeq reference GCF_000006765.1.

To determine possible reasons for the lack of mutations found in some of the phenotypic variants, the unmapped reads from the variant calling analysis were used for *de novo* assembly and annotation using SPAdes v3.10.1 or higher (*12*) and Prokka v1.12 or higher (*13*). Once it was determined that unmapped reads were likely affecting the phenotype, all sequenced isolates were assembled with SPAdes and the pangenome was visualized using Anvi'o version 6.2 (*14*, *15*) and its dependencies (*16*, *17*).

Because it appeared that exogenous DNA was inserting into the PAO1 phenotypic variants, we used MGEFinder to predict insertion loci and sequences (*18*). In addition, long read sequencing was performed to resolve the location of these insertions. DNA was prepared for the MinION (FLO-MIN106, Oxford Nanopore) using the 1D ligation SQK-LSK108 kit with native barcoding. Bases were called using the ONT Albacore Sequencing Pipeline Software (version 2.2.5). We then used a combination of the Nanopore and Illumina reads for a hybrid assembly using Unicycler v0.4.4 (*19*). Prophage regions were detected with PHASTER (*20*).

***Data accessibility***

All Illumina and Nanopore sequences have been deposited in the NCBI Sequence Read Archive (SRA) to the BioProject PRJNA633671 under accession numbers SAMN14968233 to SAMN14968292. Code for sequence processing can be found here: https://github.com/sirmicrobe/pig_wound_manuscripts/

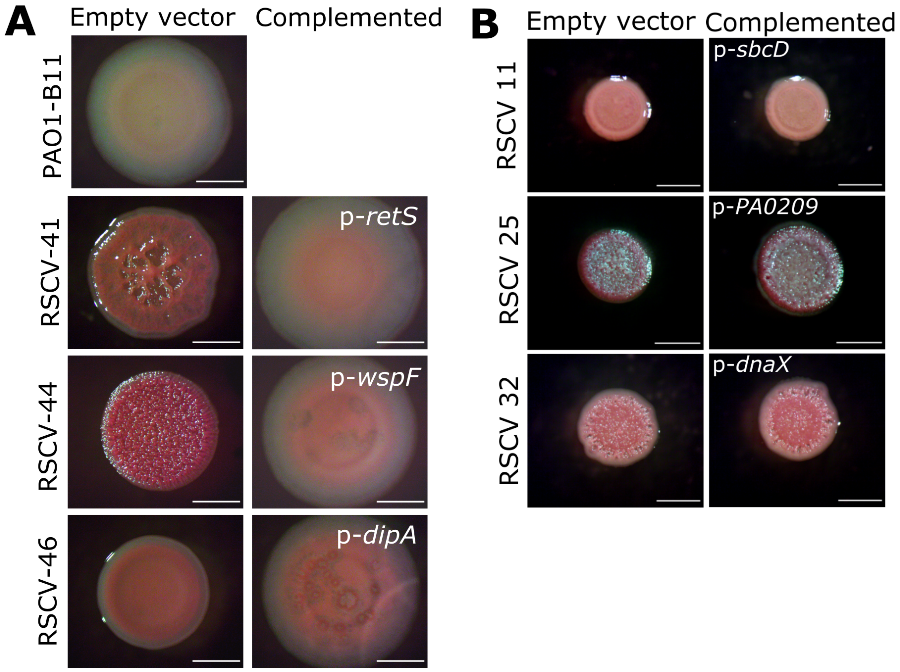


**Figure S1. Complementation of RSCVs.** PAO1 RSCVs were complemented by introducing the ancestral copy of the indicated gene. Parent strains (empty vector; pUCP18) and complemented strains were grown on VBMM and colony morphology assessed. **(A)** RSCV-41 and -44 have acquired SNPs in *retS* and *wspF* respectively. Both RSCVs were complemented by the wild type gene, indicating that these mutations are responsible for RSCV phenotype. RSCV-46 contains prophage disruptions of both *dipA* and *fapE* (amyloid synthesis). RSCV-46 was complemented by wild type *dipA*, confirming that inactivation of *dipA* was responsible for the RSCV of this isolate. **(B)** RSCV-11, -25 and -32 acquired SNPs in indicated genes that were not complemented by the ancestral copy, indicating that these SNPs are secondary to other mutations responsible for the RSCV phenotype. Scale bar indicates 2mm.

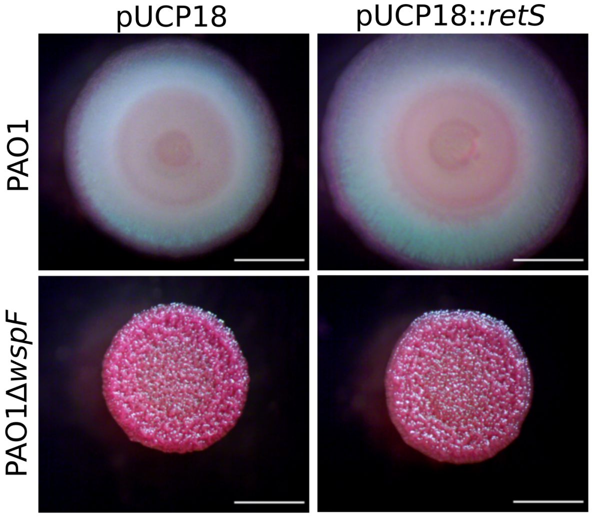


**Figure S2. Ancestral *retS* allele does not complement the RSCV phenotype in a non-specific manner.** *retS* complementing vector was introduced into PAO1Δ*wspF* and the isogenic PAO1 parent. Colony morphology was assessed on VBMM. The RSCV phenotype of PAO1Δ*wspF* was not complemented by wild type *retS*, indicating that overexpression of the global regulator, *retS*, on a high copy plasmid does not complement the general RSCV phenotype of non-*retS* mutants. Scale bar indicates 2mm.

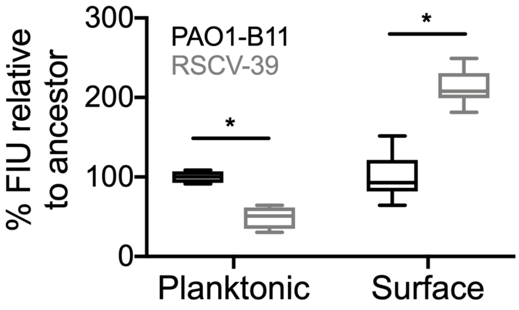


**Figure S3. RSCV-39 only displays elevated cyclic-di-GMP levels when grown on a surface**. A cyclic-di-GMP reporter was electroporated into RSCV-39 (grey) and the ancestor PAO1 (black). Strains were grown either in planktonic culture, or on solidified media and cyclic-di-GMP levels were measured as green fluorescence. Data is presented as GFP fluorescence intensity units (FIU) normalized to the ancestor strain for each growth condition, which was set at 100%. FIU was normalized to optical density. N=3, each with four technical replicates. * indicates $p<0.05$ (Student's t-test).

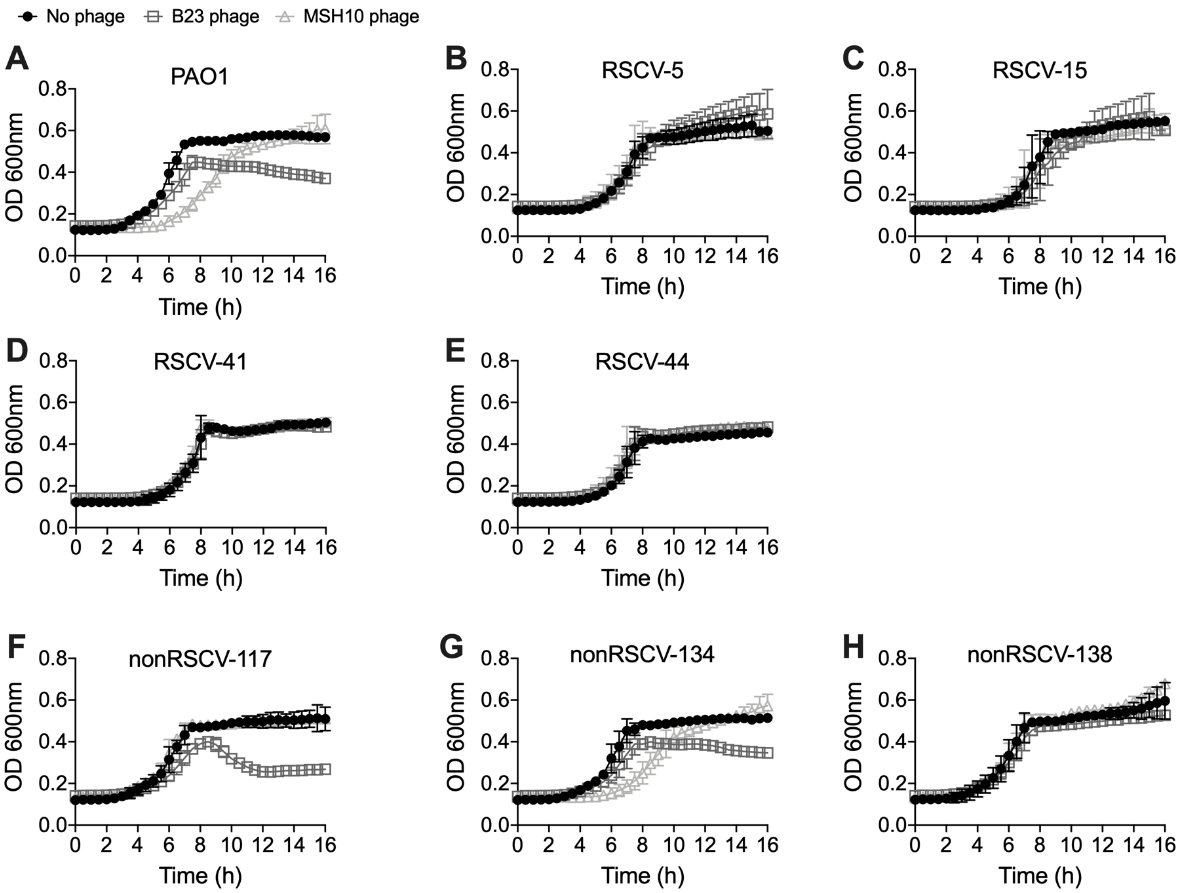


**Figure S4. PAO1 wound isolates are immune to phage infection. (A)** The ancestral PAO1 and representative **(B-E)** PAO1 RSCV and **(F-H)** nonRSCV wound isolates were grown in planktonic culture with phage isolated from B23 or MSH10 for 16 h and the OD 600 measured every 30 min. Area under the curve analysis is presented in Figure 5. N=3, each with three technical replicates. Data presented as mean ± SD.

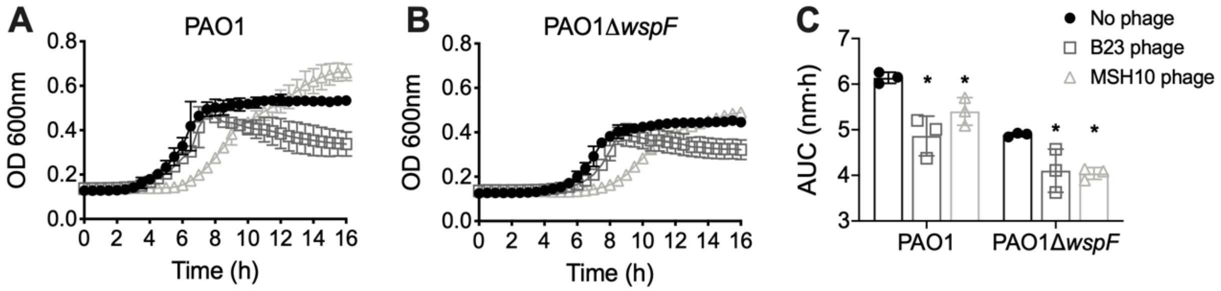


**Figure S5. PAO1Δ*wspF* is sensitive to phage infection. (A)** The parent PAO1 and **(B)** isogenic PAO1Δ*wspF* were grown in planktonic culture with phage isolated from B23 or MSH10 for 16 h and the OD 600 measured every 30 min. Data presented as mean ± SD. **(C)** Area under the curve analysis of the growth curves in **(A)** and **(B)**. * indicates p<0.05 compared to no phage control (one-way ANOVA with a Tukey's post-hoc test). Data presented as mean ± SD, with the individual data points reflecting the mean of each biological replicate. N=3, each with three technical replicates.

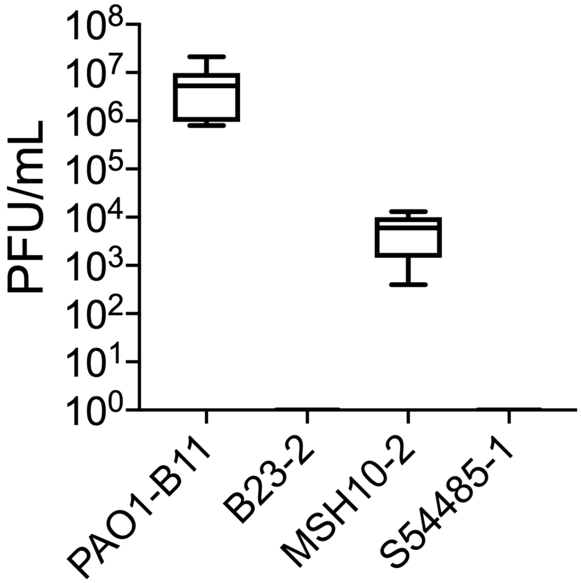


**Figure S6. Newly acquired prophage in RSCV-5 can be induced and are infectious.** Phages from RSCV-5 were isolated, and plaque assays were performed on ancestral *P. aeruginosa* strains used in the original infection, to determine if induced phages are still infectious. The PAO1-B11 ancestor and MSH10-2 were susceptible to phage infection. However, B23-2 and S54485-1 were resistant, likely due to the high similarity of RSCV-5 prophages to prophages in B23-2 and S54485-1.

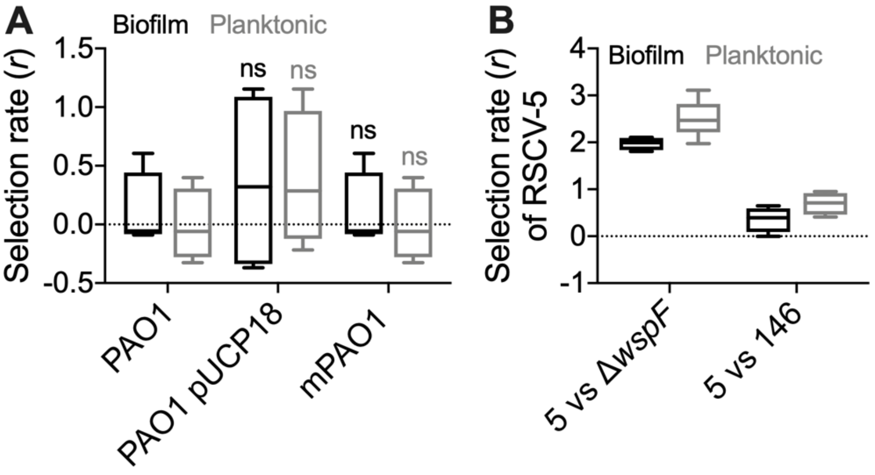


**Figure S7. Selection rate of controls** of control strains versus PAO1+ (A) and competitions between mutants (B). No significant difference between any of the strains and PAO1 vs. PAO1+ (ANOVA, $p>0.05$). N=4.

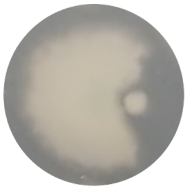

**Figure S8.** RSCV-5 colony (right) inhibiting growth of ancestral PAO1 colony (left) on an agar plate.

### Supplementary Tables

### Table S1. Mutations in PAO1 RSCV isolates

| Day | Wound # | Isolate # | Gene | Mutation |
|---|---|---|---|---|
| 14 | 4 | 5 | - | - |
| | | 11 | *sbcD* | T14P (ACC>CCC) |
| | | 15 | - | - |
| | | 18 | - | - |
| | | 19 | - | - |
| | | 46 | - | - |
| | 1 | 21 | - | - |
| | | 22 | - | - |
| | | 25 | *PA0209* | Δ55bp (721-775/882nt) |
| | | 26 | - | - |
| | | 30 | - | - |
| | | 31 | - | - |
| | | 32 | *dnaX* | +CGAGCC (1406/2046 nt) |
| | | | *PA5201* | T139P (ACC>CCC) |
| | | 34 | - | - |
| | 3 | 33 | - | - |
| | | 35 | - | - |
| | | 39 | - | - |
| 28 | 2 | 41 | *retS* | T443P(ACC>CCC) |
| | 3 | 44 | *wspF* | I68S(ATC>AGC) |
| | | | *pilQ* | +GT (656/2145nt) |
| | | | *PA2942/PA2943* | C>T intergenic (544/27) |

nt; nucleotide

**Table S2. Mutations in PAO1 non-RSCV isolates**

| Day | Wound # | Isolate # | Gene | Mutation |
|---|---|---|---|---|
| 3 | 2 | 106 | - | - |
| | 3 | 110 | - | - |
| | | 111 | - | - |
| 14 | 1 | 117 | - | - |
| | 2 | 122 | - | - |
| | | 123 | - | - |
| | | 124 | - | - |
| | | 125 | - | - |
| | | 126 | - | - |
| | 3 | 130 | - | - |
| | | 131 | - | - |
| | 4 | 134 | *PA0727* | H180H (CAT>CAC) |
| | | 136 | - | - |
| 28 | 1 | 137 | - | - |
| | | 138 | - | - |
| | | 139 | - | - |
| | | 140 | - | - |
| | 2 | 142 | *PA1680* | L46L (CTG>CTA) |
| | | 143 | *PA0788* | R472R (CGC>CGA) |
| | | 144 | - | - |
| | | 145 | *PA0065* | I26T (ATC>ACC) |
| | | | *[rbsK]* | Δ294bp (688/926nt) |
| | | 146 | - | - |
| | 3 | 149 | *pilQ* | +GT (656/2145 nt) |
| | | | *rho* | S280C (AGC>TGC) |
| | 4 | 152 | *mexT* | D67E (GAC>GAA) |
| | | 153 | *mexT* | D67E (GAC>GAA) |
| | | 154 | *mexT* | D67E (GAC>GAA) |
| | | 155 | *mexT* | D67E (GAC>GAA) |
| | | 156 | *mexT* | D67E (GAC>GAA) |

**Table S3. Number of isolates containing sequences from extinct strains**

| Isolate | Total sequenced | B23 | | MSH10 | | S54485 | |
|---|---|---|---|---|---|---|---|
| | | # | % | # | % | # | % |
| RSCV | 17 | 17 | 100 | 14 | 82 | 17 | 100 |
| non-RSCV | 28 | 14 | 50 | 18 | 64 | 14 | 50 |

**Table S5. Summary of *retS* mutations**

| Day | Wound # | Isolate # | *retS* (PA4856) mutation |
|---|---|---|---|
| 14 | 4 | 5 | Insertion at 2369 [A] |
| | | 11 | - |
| | | 15 | - |
| | | 18 | - |
| | | 19 | - |
| | | 46 | - |
| | 1 | 21 | Insertion at 2525 |
| | | 22 | Insertion at 2525 |
| | | 25 | Insertion at 2525 |
| | | 26 | Insertion at 2525 |
| | | 30 | Insertion at 1067/1780 |
| | | 31 | Insertion at 1067/1780 |
| | | 32 | Insertion at 1067/1780 |
| | | 34 | Insertion at 1067/1780 |
| | 3 | 33 | - |
| | | 35 | - |
| | | 39 | - |
| 28 | 2 | 41 | SNP at T443P |
| | 3 | 44 | - |

[A] Insertion sites are indicated by nucleotide number

**Table S6. Mutations and insertions of mobile genetic elements in PAO1 non-RSCV isolates**

| Day | Wound # | Isolate # | Gene | Mutation/ phage | S54485 plasmid |
|---|---|---|---|---|---|
| 3 | 2 | 106 | - | - | - |
| | 3 | 110 | - | - | - |
| | | 111 | - | - | - |
| 14 | 1 | 117 | *tRNA-leu (PA1796)* | F116 | + |
| | | 118 | *tRNA-leu (PA1796)* | F116 | - |
| | | 119 | *tRNA-leu (PA1796)* | F116 | - |
| | | | *PA3825* | F10 | |
| | | | *dipA* | JBD24 | |
| | | | *PA0461* | JBD24 | |
| | | 120 | *tRNA-leu (PA1796)* | F116 | - |
| | | | *PA3969a/ amn* | G | |
| | | 121 | *tRNA-leu (PA1796)* | F116 | - |
| | | | *lysR* | PA14_35710 | |
| | | | *MFS transporter (PA5311)* | G198G | |
| | 2 | 122 | *tRNA-leu (PA1796)* | F116 | - |
| | | | *lysR* | PA14_35710 | |
| | | 123 | *tRNA-leu (PA1796)* | F116 | - |
| | | 124 | *tRNA-leu (PA1796)* | F116 | - |
| | | | *lysR* | PA14_35710 | |
| | | 125 | *tRNA-leu (PA1796)* | F116 | - |
| | | 126 | *tRNA-leu (PA1796)* | F116 | - |
| | | | *retS* | JBD24 | |
| | 3 | 130 | *tRNA-leu (PA1796)* | F116 | + |
| | | | *PA3825/* | F10 | |
| | | | *PA0146* | JBD24 | |
| | | | *PA2190* | JBD24 | |
| | | | *PA3464* | JBD24 | |
| | | 131 | *tRNA-leu (PA1796)* | F116 | + |
| | | | *PA3825* | F10 | |
| | | | *PA0146* | JBD24 | |
| | | | *PA2190* | JBD24 | |
| | | | *PA3464* | JBD24 | |
| | 4 | 134 | *PA0727* | H180H | - |
| | | 136 | *tRNA-leu (PA1796)* | F116 | + |
| 28 | 1 | 137 | *tRNA-leu (PA1796)* | F116 | + |
| | | | *PA3825* | F10 | |

| | | | | | |
|---|---|---|---|---|---|
| | | | *PA2035* | JBD24 | |
| | | | *PA2318i* | JBD24 | |
| | | | *mexT* | JBD24 | |
| | | | *PA2727* | JBD24 | |
| | | 138 | *tRNA-leu (PA1796)* | F116 | + |
| | | | *PA3825* | F10 | |
| | | | *PA2035* | JBD24 | |
| | | | *mexT* | JBD24 | |
| | | | *PA2727* | JBD24 | |
| | | | *fptA* | JBD24 | |
| | | 139 | *tRNA-leu (PA1796)* | F116 | + |
| | | | *PA3825* | F10 | |
| | | | *PA2035* | JBD24 | |
| | | | *mexT* | JBD24 | |
| | | | *PA2727* | JBD24 | |
| | | | *fptA* | JBD24 | |
| | | 140 | *tRNA-leu (PA1796)* | F116 | + |
| | | | *PA3825* | F10 | |
| | | | *PA2035* | JBD24 | |
| | | | *mexT* | JBD24 | |
| | | | *eddB* | JBD24 | |
| | 2 | 142 | *tRNA-leu (PA1796)* | F116 | + |
| | | | *PA3825* | F10 | |
| | | | *PA0461* | JBD24 | |
| | | | *pauA4* | JBD24 | |
| | | | *mexT* | JBD24 | |
| | | | *PA1680* | L46L | |
| | | 143 | *tRNA-leu (PA1796)* | F116 | + |
| | | | *PA3825* | F10 | |
| | | | *PA0461* | JBD24 | |
| | | | *pauA4* | JBD24 | |
| | | | *mexT* | JBD24 | |
| | | | *PA0788* | R472R | |
| | | 144 | *tRNA-leu (PA1796)* | F116 | + |
| | | | *PA3825* | F10 | |
| | | | *PA0461* | JBD24 | |
| | | | *pauA4* | JBD24 | |
| | | | *mexT* | JBD24 | |
| | | 145 | *tRNA-leu (PA1796)* | F116 | + |

| | | | | |
|---|---|---|---|---|
| | | *lysR* | PA14_35710 | |
| | | *PA3825* | F10 | |
| | | *phzS* | JBD24 | |
| | | *PA4453* | JBD24 | |
| | | *PA0065* | I26T | |
| | | *[rbsK]* | Δ294bp | |
| | 146 | *tRNA-leu (PA1796)* | F116 | + |
| | | *lysR* | PA14_35710 | |
| 3 | 149 | *pilQ* | +GT | + |
| | | *rho* | C280S | |
| 4 | 152 | *PA3457i* | PA0445 transposase | - |
| | | *mexT* | D67E | |
| | 153 | *PA3457i* | PA0445 transposase | - |
| | | *mexT* | D67E | |
| | 154 | *PA3457i* | PA0445 transposase | - |
| | | *mexT* | D67E | |
| | 155 | *PA3457i* | PA0445 transposase | - |
| | | *mexT* | D67E | |
| | 156 | *PA3457i* | PA0445 transposase | - |
| | | *mexT* | D67E | |

**Table S7: Strains and plasmids used in this study**

| Strain or plasmid | Relevant genotype/characteristics | Ref or source |
|---|---|---|
| *P. aeruginosa* | | |
| PA14-1 | Barcode CAAAAGGACA. Gent | Gloag *et al.* 2019 |
| PAO1-B11 | Barcode GTGTCGTGGG. Gent | Gloag *et al.* 2019 |
| B23-2 | Wound isolate. Barcode GCCTATTGTG. Gent | Gloag *et al.* 2019 |
| CF18-1 | CF isolate. Barcode GTTACGTCAA. Gent | Gloag *et al.* 2019 |
| MSH10-2 | Water isolate. Barcode TATCAGATTT. Gent | Gloag *et al.* 2019 |
| S54485-1 | UTI isolate. Barcode TTAAACTAGG. Gent | Gloag *et al.* 2019 |
| PAO1 | Wild type parent of *wspF* deletion | |
| PAO1Δ*wspF* | Clean *wspF* deletion (JJH356) | |
| mPAO1 | Wild type parent of transpson library | Jacobs *et al.* 2003 |
| mPAO1*retS*::Tn | Transposon insertion in *retS* (PW9164) | Jacobs *et al.* 2003 |
| PAO1*attB*::*lacZ* | *lacZ* from miniCTX-*lacZ* introduced at the *attB* site | This study |
| PAO1 pUCP18 | Wild type PAO1 with pUCP18 empty vector | This study |
| RSCV-5 pUPC18::*retS* | RSCV-5 complemented with wild type allele of *retS* | This study |
| *E. coli* | | |
| NEB5-α | | NEB |
| S17 | | NEB |
| Plasmids | | |
| pCdrA::*gfp* | CdrA promoter fused to *gfp.* Carb | Rybtke *et al.* 2012 |
| pMH487 | Empty vector for pCdrA::*gfp.* Carb | Rybtke *et al.* 2012 |
| pUCP18 | Empty vector for complementing constructs. Carb | |
| pUCP18::retS | Complementing vector. Wild type allele amplified from PAO1 as EcoRI and HIndIII fragment. Carb | This study |
| pUCP18::*dipA* | Complementing vector. Wild type allele amplified from PAO1 as EcoRI and HIndIII fragment. Carb | This study |
| pUCP18::*dnaX* | Complementing vector. Wild type allele amplified from PAO1 as EcoRI and HIndIII fragment. Carb | This study |
| pUCP18::*sbcD* | Complementing vector. Wild type allele amplified from PAO1 as EcoRI and HIndIII fragment. Carb | This study |
| pUPC18::*PA0209* | Complementing vector. Wild type allele amplified from PAO1 as EcoRI and HIndIII fragment. Carb | This study |
| pSP5 | Complementing vector. Wild type *wspF* cloned into pUCP18. Carb | Starkey *et al.* 2009 |
| miniCTX-*lacZ* | Tet | Hoang *et al.* 2000 |

**Table S8. Primers used in this study.**

| Primer name | Primer sequence (5'-3')[A] | RE[B] |
|---|---|---|
| *retS*_F | AACCGGAATTCCCACGGCCACTTGGCTATAA | EcoRI |
| *retS*_R | CCCAAGCTTTAGCCGCGTGCGGTTATC | HindIII |
| *dipA*_F | AACCGGAATTCCTCGCTCAGTACCTGGAATCA | EcoRI |
| *dipA*_R | CCCAAGCTTCTCAGGGGGTAGGCGAATG | HindIII |
| *dnaX*_F | AACCGGAATTCCATTGGCCGACTTGTGCTTC | EcoRI |
| *dnaX*_R | CCCAAGCTTCGAATGTCGTCTTGTTGCGT | HindIII |
| *sbcD*_F | AACCGGAATTCATAAGGAATGCGAGCCACGG | EcoRI |
| *sbcD*_R | CCCAAGCTTCGAGGTTCTTCAGGCGGATG | HindIII |
| *PA0209*_F | AACCGGAATTCGAGCTGGTCGAGTGGTCC | EcoRI |
| *PA0209*_R | CCCAAGCTTTTCGAAGGTCAGGGTTTCCA | HindIII |

[A] Red indicates the restriction enzyme sequence

[B] RE indicated restriction enzyme

**Movie S1-S4.**

Colony competitions between ancestor and ancestor (1), ancestor PAO1 and RSCV-5 retS phage mutant (2), ancestor PAO1 and RSCV-41 retS SNP mutant (3), and ancestor PAO1 and RSCV-15 dipA mutant (4). The ancestral PAO1 strain is on the left in each movie.

.
